# Mis6/CENP-I maintains CENP-A nucleosomes against centromeric non-coding transcription during mitosis

**DOI:** 10.1101/2021.11.03.466203

**Authors:** Hayato Hirai, Yuki Shogaki, Masamitsu Sato

## Abstract

Centromeres are established by nucleosomes containing the histone H3 variant CENP-A. CENP-A is recruited to centromeres by the Mis18–HJURP machinery. During mitosis, CENP-A recruitment ceases, implying the necessity of CENP-A maintenance at centromeres, although the exact underlying mechanism remains elusive. Herein, we show that the kinetochore protein Mis6 (CENP-I) retains CENP-A during mitosis in fission yeast. Eliminating Mis6 during mitosis caused immediate loss of pre-existing CENP-A at centromeres. CENP-A loss occurred due to the transcriptional upregulation of non-coding RNAs at the central core region of centromeres, as confirmed by the observation RNA polymerase II inhibition preventing CENP-A loss from centromeres in the *mis6* mutant. Thus, we concluded that Mis6 blocks the indiscriminate transcription of non-coding RNAs at the core centromere, thereby retaining the epigenetic inheritance of CENP-A during mitosis.

## Introduction

Duplicated chromosomes must be equally distributed between daughter cells so that genetic information is adequately propagated to the progeny. The mitotic spindle attaches to the kinetochore, which is a macromolecular protein complex formed at the centromere, in order to equally separate sister chromatids to opposite spindle poles^1^.

In most eukaryotes, kinetochore assembly requires the deposition of CENP-A (Cnp1 in the fission yeast *Schizosaccharomyces pombe*), a centromere-specific variant of histone H3. In budding yeast, the centromere is composed of a 125-bp DNA sequence (called a point centromere), and a single histone octamer containing CENP-A is allocated to the centromere in addition to the usual H3-containing nucleosomes^2^. In contrast, multiple CENP-A-containing nucleosomes are deposited to the regional centromere, which consists of a 35∼110-kb central core region flanked by pericentric regions in fission yeast^3^, or megabases of repetitive DNA sequences in higher eukaryotes, including humans^4^.

Reduction of CENP-A levels at the centromere causes errors in chromosome segregation, which may result in aneuploidy, leading to cell death and birth defects^5^. Thus, the amount of CENP-A at the centromere is strictly regulated. Upon DNA replication, the number of nucleosomes containing CENP-A at the centromere is halved^6^, necessitating the replenishment of new CENP-A nucleosomes during the cell cycle especially prior to mitosis onset in order to assemble functional kinetochores. The timing of CENP-A supply differs between organisms, occurring in the early G1 phase in humans as opposed to mainly in G2 in fission yeast^6,7^.

Recruitment of CENP-A requires the Mis18 complex (Mis18BP1, Mis18α and Mis18β in humans^8^; Mis16, Mis18 and Mis19 (also known as Eic1 or Kis1) in fission yeast^9-12^) as well as a chaperone HJURP (Holliday junction recognition protein; Scm3 in fission yeast)^13-16^. In budding yeast^17^, human cells^18^ and *Drosophila* cells^19^, histone H3 within euchromatin undergoes dynamic turnover during the cell cycle, while CENP-A nucleosomes are generally immobile. Once recruited to centromeres, they appear neither disassembled nor replaced by new nucleosomes containing CENP-A or H3 during both mitotic cycles and meiosis in higher eukaryotes^6,20^, implying the existence of machinery for CENP-A maintenance.

In various organisms, non-coding RNAs (ncRNAs) are transcribed at the central core region of centromeres^21^. When RNA polymerase II (RNAPII) progresses to the central core region of centromeric DNA to transcribe ncRNAs, nucleosomes are temporarily evicted from the DNA. Transcription-coupled turnover of CENP-A has also been reported when transcription of the alphoid array is artificially enhanced by tethering the herpes virus VP16 activation domain to a human artificial chromosome (HAC)^22^. Furthermore, *S. pombe* Cnp1 (CENP-A) dispersed when ncRNA transcription at the central core region was upregulated in cells lacking Mediator complex subunit Med20^23^. These findings indicate that CENP-A needs to be maintained against the transcription of centromeric ncRNAs, even outside DNA replication.

While CENP-A maintenance at centromeres is necessary during the cell cycle, the underlying molecular mechanism remains elusive. Ubiquitylation of CENP-A was recently shown to contribute to CENP-A maintenance in human cells^24^. Further, human kinetochore proteins CENP-C and CENP-N directly interact with CENP-A-containing nucleosomes in *vitro*^25,26^. These interactions are required for the immobility of CENP-A-containing nucleosomes in human cells^27,28^. Another study suggested that CENP-C and CENP-N do not contribute to CENP-A maintenance in human cells^29^. Taken together, whether these factors are required for CENP-A retention in *vivo* remains controversial. It was recently demonstrated that HJURP is necessary for CENP-A maintenance during DNA replication^30^. However, HJURP dissociates from centromeres in metaphase both in human cells^8,13,14^ and in fission yeast^9,15,16^, indicating that HJURP does not engage in CENP-A maintenance during metaphase. It was recently reported that the histone chaperone Spt6, which is known as a histone H3 recycler, also contributes to the recycling of pre-existing CENP-A during ncRNA transcription at centromeres in both *Drosophila* and human cells^31^. However, whether more factors are involved in CENP-A maintenance remains unclear.

In this study, we demonstrate that the kinetochore protein Mis6 (CENP-I in human), which has been originally defined as a loading factor for CENP-A during interphase, contributes to maintenance of CENP-A during metaphase. We propose that Mis6 may counteract progression of RNAPII into core centromeres to prevent the reduction of CENP-A nucleosomes during metaphase.

## Results

### Cnp1 is not recruited to centromeres during metaphase

The Mis18 complex and Scm3 (HJURP) are loading factors for Cnp1 (CENP-A) conserved in fission yeast and higher eukaryotes, such as chickens and humans^8,9,13-16^. In fission yeast, the kinetochore protein Mis6 (CENP-I) is considered another loading factor, as Cnp1 localisation is reduced in the *mis6-302* temperature-sensitive mutant^32^. The Mis18 complex and Scm3 localise to kinetochores during interphase but are dispersed during mitosis, whereas Mis6 remains throughout the cell cycle^9,15,16,33^. This suggests that Cnp1 could be recruited at any time, assuming that Mis6 would constantly serve as a Cnp1 loading factor.

To test this possibility, we examined the kinetics of Cnp1 localisation to centromeres using Cnp1-GFP (green fluorescent protein was tagged to the C-terminus of Cnp1) strains. Cnp1-GFP cells were arrested at G1, G2 and metaphase via *cdc10-27, cdc25-22*^34^ and *alp12-1828*^35^ mutations, respectively. We then performed fluorescence recovery after photobleaching (FRAP) assays, where the Cnp1-GFP fluorescence intensity at centromeres was measured every 5 min after laser irradiation of the Cnp1-GFP foci. We found that Cnp1-GFP fluorescence intensity gradually recovered after irradiation in G1- or G2-arrested cells (**Fig. 1a–d**), indicating that bleached Cnp1 was replaced, albeit slowly. In contrast, Cnp1-GFP intensity did not recover after irradiation in metaphase-arrested cells (**Fig. 1e, f**). These results indicate that Cnp1 is not recruited to centromeres during metaphase, and Mis6 does not play a role in Cnp1 loading in the meantime, whereas Mis6 does contribute to Cnp1 recruitment during interphase by allocating Scm3 to centromeres^16^.

**Fig. 1.**
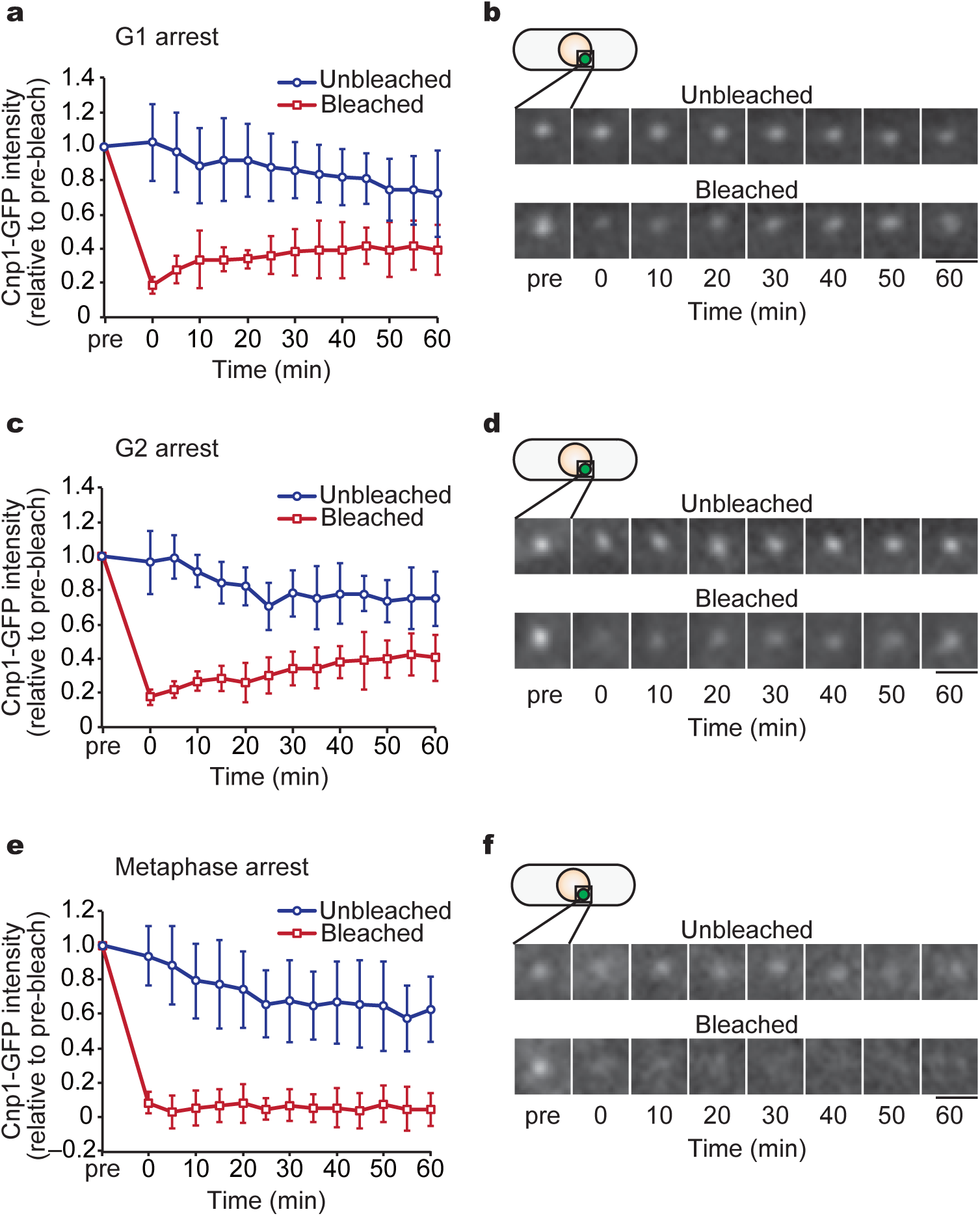
Deposition of Cnp1 (CENP-A) ceases in metaphase. Fluorescence recovery after photobleaching (FRAP) assays were performed with Cnp1-GFP cells arrested at each stage of the cell cycle. Cells with unbleached centromeres were used as controls. **(a**,**c**,**e)** Recovery kinetics of Cnp1-GFP fluorescence over time. Intensities are shown normalised to values before bleaching (pre). **(b**,**d**,**f)** Time-lapse Cnp1-GFP images of representative cells. Cells were arrested in G1 (using the *cdc10* mutant; a,b), G2 (*cdc25*; c,d) or metaphase (*alp12*; e,f), and Cnp1-GFP dots were photobleached. Fluorescence recovery at the centromeres was then monitored over time. Numbers of unbleached and bleached cells: *n* = 11 and 16 (a); *n* = 9 and 9 (c); *n* = 19 and 15 (e), respectively. Data are presented as the mean ± s.d. Scale bars = 2 *µ*m.

### Mis6 is required to maintain Cnp1 at centromeres during metaphase

We speculated that Mis6 may play another role at the centromeres during metaphase. We first examined whether the function of Mis6 during metaphase is essential for viability. The *mis6-302* mutant was combined with the *cut9-665* mutant, which has defects in cell cycle transition from metaphase to anaphase because of the low APC/C (anaphase promoting complex/cyclosome) activity at restrictive temperature^36^. The resultant *mis6-302 cut9-665* double mutant exhibited severe growth defects, even at semi-restrictive temperature (**Fig. 2a**). This suggests that the mitotic function of Mis6 is crucial for prolonged metaphase.

**Fig. 2.**
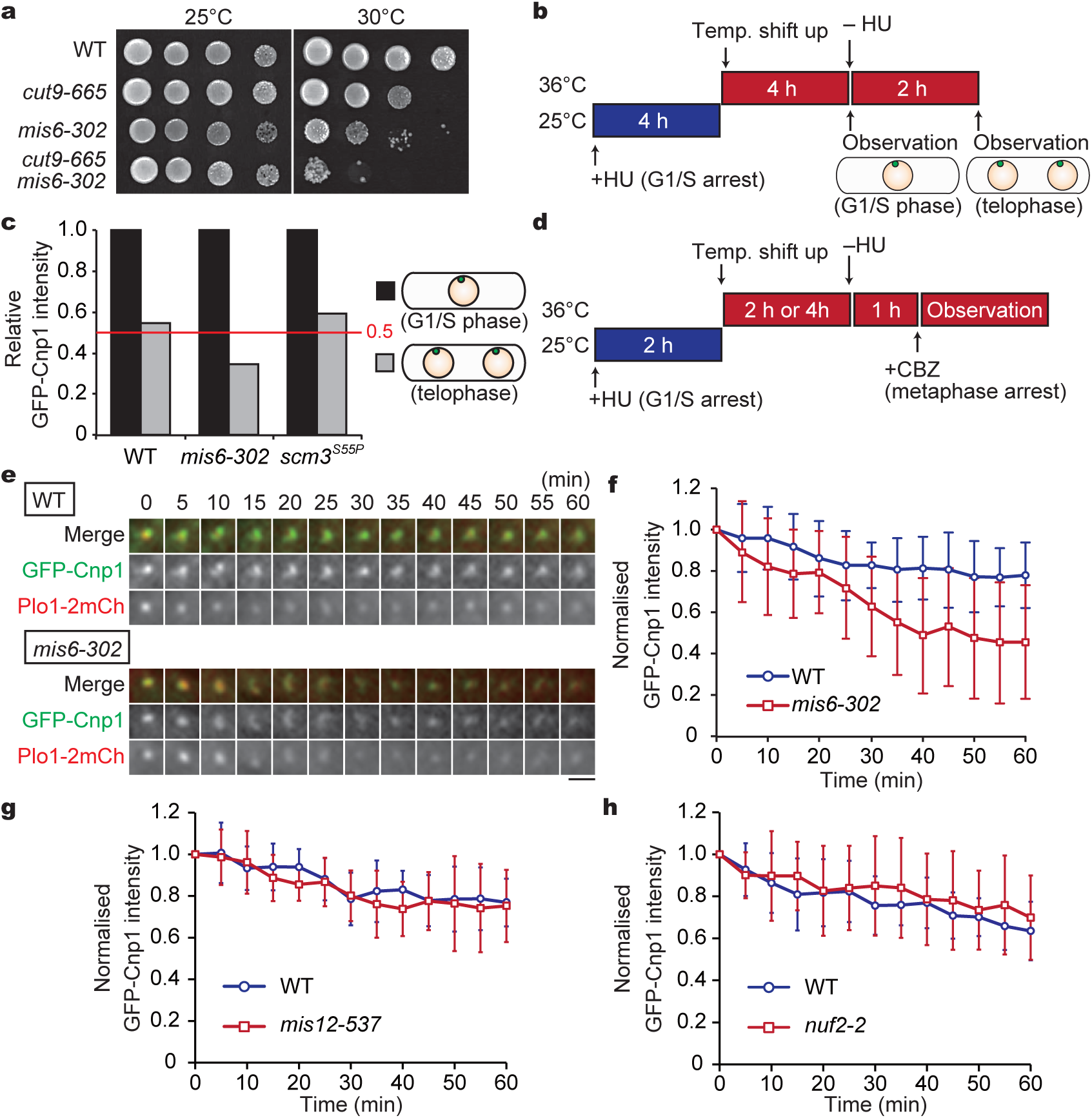
Mis6 is required for Cnp1 maintenance at the centromere during metaphase. **(a)** Growth assays for indicated strains. Cells at 10-fold serial dilutions were grown at 25 and 30°C. **(b**,**c)** Comparison of GFP-Cnp1 signals before and after mitosis. Experimental design (b). Cells were synchronised at G1/S using hydroxyurea (HU) at 25°C, followed by a shift up to the restrictive temperature (36°C) for *mis6-302* and *scm3*^*S55P*^ mutants in advance. The cells were then released to late mitosis via HU removal. GFP-Cnp1 fluorescence intensity was measured before (G1/S) and after release from HU (telophase). GFP-Cnp1 intensities in WT, *mis6-302* and *scm3*^*S55P*^ cells were quantified (*n* ≥ 22 cells, c). The data is normalised as a ratio of values for telophase to interphase. **(d–h)** Cnp1 maintenance assays using GFP-Cnp1 Plo1-2mCherry cells of the WT, *mis6-302, mis12-537* and *nuf2-2* background. **(d)** Experimental design. Cells were synchronised at G1/S via HU treatment at 25°C and shifted up to 36°C for 2 h to inactivate Mis6 and Alp12 or for 4 h to inactivate Mis12, Nuf2 and Alp12 in advance. Cells were released from HU and arrested in mitosis using CBZ. **(e)** Time-lapse images of the GFP-Cnp1 signal (green) during metaphase in WT and *mis6-302* cells. Plo1-2mCh (red) is shown as a mitotic marker (Scale bar: 2 *µ*m). **(f–h)** Temporal kinetics of GFP-Cnp1 intensities during metaphase arrest, normalised to values for 0 min. **(f)** WT, *n* = 16 cells; *mis6-302, n* = 20 cells. Data are presented as the mean ± s.d. **(g)** WT, *n* = 11 cells; *mis12-537, n* = 6 cells. Data are presented as the mean ± s.d. **(h)** WT, *n* = 21 cells; *nuf2-2, n* = 18 cells. Data are presented as the mean ± s.d.

We then tried to determine the function of Mis6 during metaphase. As overexpression of Cnp1 suppressed the temperature sensitivity of the *mis6-302* mutant, albeit partially^9,37^, the mitotic function of Mis6 would also be exerted for positive regulation of Cnp1. We then predicted that Mis6 plays a role in the maintenance of Cnp1 during metaphase.

To test this hypothesis, we performed assays to monitor Cnp1 maintenance during metaphase: WT (*mis6*^+^) and *mis6-302* cells expressing GFP-Cnp1 were synchronously arrested in metaphase in order to follow the kinetics of Cnp1 localisation to centromeres during metaphase. As illustrated in **Fig. 2b**, cells were first arrested in the G1/S phase using hydroxyurea (HU), followed by a temperature shift up to 36°C in order to inactivate Mis6 before mitotic entry. After HU washout, cells were released into mitosis until telophase (mitotic exit).

In WT cells, the intensity of a single GFP-Cnp1 dot in telophase was approximately half that in the previous G1/S stage. This reduction in GFP-Cnp1 intensity simply reflects the segregation of sister chromatids in the meantime. Therefore, Cnp1 was rarely removed from the centromeres during WT mitosis. This result is consistent with previous observations in human cells showing that pre-existing CENP-A at centromeres is retained throughout the cell cycle^6^. In the *mis6-302* mutant, GFP-Cnp1 intensity during telophase was approximately 35% of that in G1/S (**Fig. 2c**), indicating that Cnp1 was dissociated from centromeres when cells passed through mitosis in the absence of Mis6. Thus, Mis6 is required for Cnp1 retention during mitosis.

It could be hypothesised that Scm3 (HJURP), rather than Mis6, is responsible for Cnp1 maintenance, because Scm3 and Mis6 are interdependent for localisation to centromeres^15^. This is unlikely, as Scm3 ceases localisation to centromeres in WT cells during mitosis^15,16^. To further clarify this point, we isolated a temperature-sensitive *scm3* mutant, in which the localisation of Mis6-GFP was unaffected at restrictive temperature (**Supplementary Fig. S1 a–c**). The mutant harboured a substitution at position 55 (*scm3*^*S55P*^), which was close to that in the previous *scm3*^*L56F*^ mutant that also retained Mis6 localisation to centromeres^16^. In the *scm3*^*S55P*^ mutant, the reduction in GFP-Cnp1 intensity after mitosis was almost comparable to that in WT cells (∼60 %, **Fig. 2c**), indicating that Cnp1 did not dissociate from centromeres during metaphase. Thus, Scm3 was dispensable for the maintenance of mitotic Cnp1.

To further investigate whether Mis6 retains Cnp1 during metaphase, WT and *mis6-302* cells expressing GFP-Cnp1 were prepared similarly as in **Fig. 2b**, but arrested to mitosis (metaphase) for ≥ 1 h by double inhibition using the microtubule poison carbendazim (CBZ) in combination with the α-tubulin temperature-sensitive mutant *alp12-1828*^35^ (**Fig. 2d**). Metaphase arrest of the cell was confirmed by observing the Plo1 (Polo-like kinase) foci at spindle pole bodies^38,39^. The GFP-Cnp1 intensity at centromeres decayed more rapidly in *mis6-302* cells than in WT cells (**Fig. 2e, f**). Taken together, these results suggest that Mis6, but not Scm3, is responsible for the maintenance of Cnp1 at centromeres during metaphase.

We examined whether the role of Cnp1 maintenance at mitotic centromeres is specifically assigned to Mis6 or shared with other kinetochore components by repeating similar GFP-Cnp1 maintenance assays using the *mis12-537* and *nuf2-2* mutants. These harbour mutations in Mis12 (belonging to the Mis12/Mtw1 subcomplex) and Nuf2 (to the Ndc80 subcomplex), respectively^40,41^. These components localised to the outer regions of centromeres relative to Mis6, and Mis6 localised to centromeres in the *mis12* and *nuf2* mutants (**Supplementary Fig. S1d**)^41,42^.

In both mutants, signal intensities of Cnp1 at centromeres during mitotic arrest were retained as in the WT and unlike in the *mis6-302* mutant (**Fig. 2g, h**). These assays indicated that involvement in Cnp1 maintenance is not a common feature among kinetochore factors. Rather, it is specifically assigned to Mis6.

### Enforced dismissal of Mis6 during metaphase reduces Cnp1

To further examine the function of Mis6 during mitosis, we sought to introduce experimental conditions in which Mis6 remains functional during interphase but is rapidly inactivated upon entry into mitosis. We developed a strategy for knocksideways experiments using Mis6-GFP and Kis1-GBP (GFP-binding protein^43^), as illustrated in **Fig. 3a**. Kis1 (Mis19 or Eic1) together with Mis16 and Mis18 forms the Mis18 complex to recruit Cnp1 to centromeres during interphase, whereas the whole complex disperses during metaphase^9-12^ (top, **Fig. 3a**). Taking advantage of the turnover, the fusion protein Kis1-GBP may dismount Mis6-GFP from centromeres upon mitotic entry (bottom, **Fig. 3a**). As shown in **Fig. 3b**, Mis6-GFP and Kis1-GBP-mCherry co-localised to centromeres during interphase. In metaphase cells, the Mis6-GFP signal at centromeres substantially decreased, indicating that Mis6-GFP was removed from centromeres specifically during mitosis by the knocksideways system, as expected. Later in anaphase, Mis6-GFP and Kis1-GBP-mCherry re-localised together at centromeres.

**Fig. 3.**
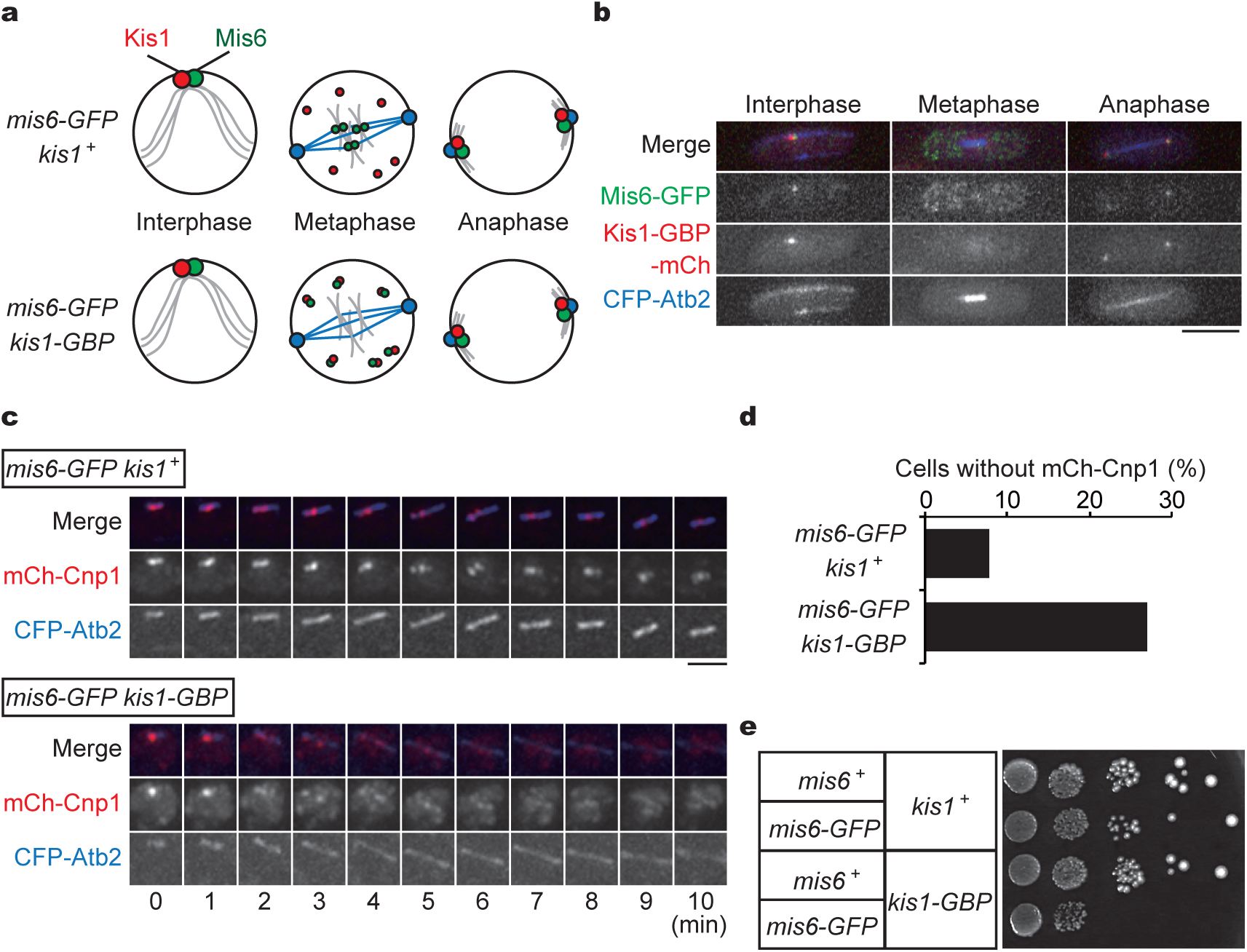
Metaphase-specific removal of Mis6 causes a decrease of Cnp1. **(a)** Schematics illustrating the outline of knocksideways to remove Mis6 from centromeres only during metaphase using Kis1. Kis1 (red) and Mis6 (green) co-localise to centromeres during interphase, but Kis1 disperses at mitosis onset (top). In cells co-expressing Kis1-GBP and Mis6-GFP, Kis1-GBP brings Mis6-GFP out of centromeres only during metaphase (bottom). **(b)** Cells expressing Mis6-GFP (green), Kis1-GBP-mCherry (red) and CFP-Atb2 (cyan fluorescent protein fused with α2-tubulin, blue) at each cell cycle stage. Scale bar = 5 *µ*m. **(c)** Pro ∼ metaphase cells expressing Mis6-GFP, mCh-Cnp1 (red) and CFP-Atb2 (microtubules; cyan) together with untagged Kis1 (top) or Kis1-GBP (bottom) were imaged. Scale bar = 2 *µ*m. **(d)** Population of metaphase cells with or without mCherry-Cnp1 at centromeres (*n* > 50 cells). **(e)** Growth assays for indicated strains. Cells at 10-fold serial dilutions were grown in EMM at 36°C.

We then evaluated whether Cnp1 localisation is reduced in response to the enforced removal of Mis6 during metaphase. In control cells (*mis6-GFP kis1*^+^, **Fig. 3c**), mCherry-Cnp1 foci at centromeres were constantly detected along the spindle during prometaphase, and only 8% of control cells failed to retain mCherry-Cnp1 at centromeres (**Fig. 3d**). In contrast, in *mis6-GFP kis1-GBP* double-tagged cells, mCherry-Cnp1 signals were clearly detected at centromeres during the initial stage of mitosis (0 min, **Fig. 3c** and **Supplementary Fig. S2**), verifying that co-expression of Mis6-GFP and Kis1-GBP did not affect their functions in Cnp1 deposition during interphase. In contrast, immediately after mitosis onset, mCherry-Cnp1 signals dispersed from centromeres in ∼25% of the cells (**Fig. 3c, d**). Moreover, the double-tagged strain displayed growth defects at high temperature, further supporting the importance of Mis6 localisation during mitosis (**Fig. 3e**).

### Reduction of Cnp1 during metaphase is coupled to transcription at centromeres

A major reason for Cnp1 loss during mitosis may be the transcription of ncRNAs in centromeres. ncRNAs are transcribed in various organisms, including plants, fission yeast and humans^21^. As our results indicate that Mis6 retains Cnp1 to centromeres, we focused on the relationship between ncRNA transcription at the central core region of centromeres and Cnp1 maintenance. The central core region (*cnt*) of centromeres is transcriptionally silenced in WT cells, whereas silencing is impaired in the *mis6-302* mutant^37^. This was reproduced in our experiments using the *cnt1::ura4*^+^ strain, in which the *ura4*^+^ reporter gene conferring the uracil autotroph was inserted in the central core region of chromosome 1 (**Fig. 4a**). Growth of WT cells harbouring the *cnt1::ura4*^+^ insertion was defective in medium lacking uracil (SD–Ura) and was fine in the counter-selection medium containing 5-fluoroorotic acid (FOA). This indicated that transcription in the central core region is subtle and is mostly silenced. In contrast, the *mis6-302* mutant containing *cnt1::ura4*^+^ showed the opposite growth pattern, confirming that the central core region in the *mis6-302* mutant was no longer silenced (**Fig. 4a**).

**Fig. 4.**
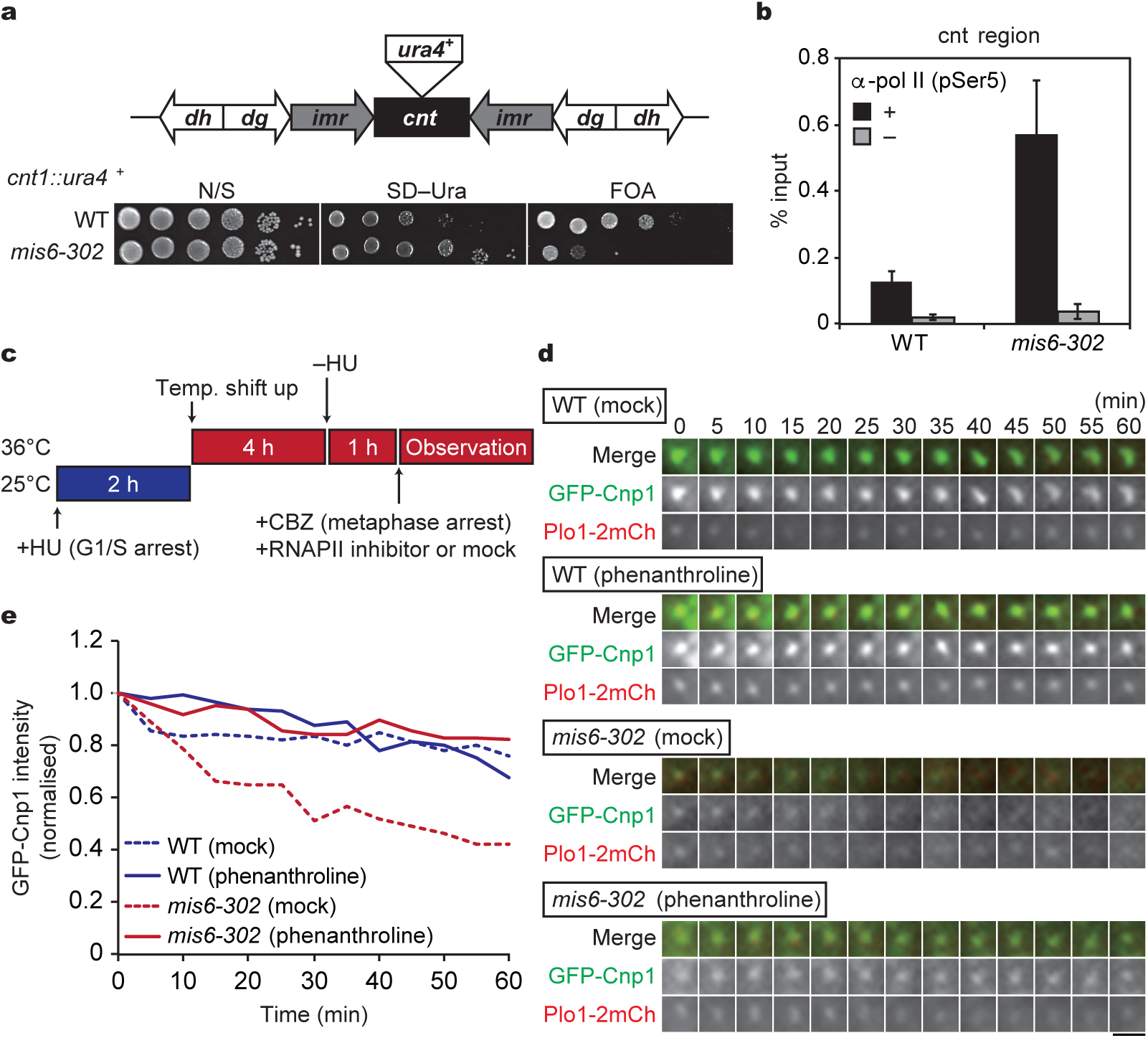
Loss of Cnp1 in *mis6-302* cells is coupled with centromeric transcription by RNA polymerase II. **(a)** A schematic illustrating the centromeric DNA structure (top). For silencing assays, the *ura4*^+^ gene is inserted in the central core (*cnt*) region (*cnt::ura4*^+^) in chromosome I. *imr*, the innermost repeats; *dg* and *dh*: the outer repeats. Silencing assays (bottom). 10-fold serial dilution of WT and *mis6-302* cells with *cnt::ura4*^+^, grown on nonselective (N/S), uracil-deficient (SD–Ura) and counter-selective (FOA) media at 25°C. **(b)** Chromatin IP (ChIP) for phosphorylated RNA polymerase II (RNAPII pSer5) at *cnt* region (the central core region) in WT and *mis6-302* cells. ChIP was performed with (+) or without (–) the anti-pol II (pSer5) antibody. Error bars = ± s. d. (*N* = 2 experiments). **(c–e)** RNAPII was inhibited in WT and *mis6-302* cells expressing GFP-Cnp1 and Plo1-2mCherry. **(c)** Experimental design. Cells arrested to G1/S at 25°C were shifted up to 36°C to inactivate Mis6 or Alp12. The cells were then released into mitosis and arrested at metaphase using CBZ. Cells were treated with an RNAPII inhibitor (1,10-phenanthroline) or mock treatment prior to observation. **(d)** Time-lapse imaging of GFP-Cnp1 (green) with Plo1-2mCh (red) in WT and *mis6-302* cells with or without the inhibitor. Scale bar = 2 *µ*m. **(e)** A Cnp1 maintenance assay. Temporal kinetics of the GFP-Cnp1 fluorescence intensity during metaphase in the indicated samples were monitored. WT (mock), *n* = 22 cells; *mis6-302* (mock), *n* = 17 cells; WT (phenanthroline), *n* = 30 cells; *mis6-302* (phenanthroline), *n* = 20 cells. The data are normalised to the intensities at 0 min, and the mean is shown.

Likewise, *mis12-537* and *nuf2-2* mutants were used in centromeric silencing assays. In contrast to the *mis6-302* mutant, both *mis12-537* and *nuf2-2* mutants showed restricted growth in the medium lacking uracil and retained their growth capacity in the presence of FOA at semi-restrictive temperatures (**Supplementary Fig. S3a**). These results demonstrate that kinetochore mutants with intact Mis6 localisation retained the ability to silence transcription at the central core region.

Chromatin IP experiments using an antibody against the C-terminal-phosphorylated form (pSer5) of RNA polymerase II (RNAPII) detected enrichment of the central core sequences in the *mis6-302* mutant compared to WT cells (**Fig. 4b**). In contrast, the enrichment was undetectable in both the *mis12-537* and *nuf2-2* mutants (**Supplementary Fig. S3b**), wherein Mis6 remained at centromeres. This demonstrates that active RNAPII transcribes ncRNA in the central core region when Mis6 is absent.

In general, conventional histone octamers containing H3 are temporarily dismantled by the chromatin remodelling factor Fun30 (Fft3 in fission yeast), allowing RNAPII to proceed with transcription^44-46^.

Analogously, Cnp1-containing octamers must be temporarily dissociated from centromeres upon transcription, which could be a potential reason for Cnp1 loss in mitotic *mis6-302* cells. It is possible that Fft3 also removes Cnp1-containing octamers upon ncRNA transcription at the centromeres. When we followed the temporal kinetics of GFP-Cnp1 intensity during metaphase, the reduction of GFP-Cnp1 intensity seen in *mis6-302* cells was cancelled by the additional knockout of Fft3 (*mis6-302 fft3Δ*, **Supplementary Fig. S4a**), confirming that Fft3 removes Cnp1 upon transcription at centromeres.

To find further evidence of transcription-coupled Cnp1 dismantling, we tested whether Cnp1 loss still occurs when transcription is blocked by RNAPII inhibitors such as 1,10-phenanthroline and thiolutin. Experiments were designed as shown in **Fig. 2d**. In brief, the cells were arrested to metaphase using CBZ as well as through *alp12-1828* mutation, and GFP-Cnp1 intensity was monitored in the presence or absence of RNAPII inhibitors (**Fig. 4c**). In WT cells, RNAPII inhibitor 1,10-phenanthroline did not affect the amount of Cnp1 at centromeres (WT, **Fig. 4d, e**). In *mis6-302* cells without RNAPII inhibitor treatment, the GFP-Cnp1 intensity constantly decreased throughout metaphase, as shown in **Fig. 2e, f**. In contrast, 1,10-phenanthroline treatment resulted in similar GFP-Cnp1 intensity as in WT cells (**Fig. 4d, e**). Comparable results were obtained in assays using thiolutin as the RNAPII inhibitor (**Supplementary Fig. S4b, c**). Taken together, the reduction of GFP-Cnp1 during mitosis of the *mis6-302* mutant is coupled to the transcription of ncRNAs at the central core region.

These results demonstrated that Mis6 prevents unnecessary transcription of ncRNAs at the central core region, thereby maintaining Cnp1 on chromatin during metaphase.

### Chromatin remodelling factor Spt6 is required for Cnp1 recycling during mitosis

Transcription of ncRNAs in the central core region occurs at a certain level even in the presence of functional Mis6, as a minor amount of phosphorylated RNAPII was detected in the central core region (see **Fig. 4b**). It has been previously shown that phosphorylated RNAPII at the central core region is upregulated during the G2-M transition^47^. Thus, some RNAPII passes through the central core region due to the removal of Cnp1 by Fft3 in WT cells, while the Cnp1 fluorescence signal remained during metaphase (**Figs. 2, 3**). These results imply that Cnp1 is maintained through additional mechanisms, other than the Mis6-dependent system, during ncRNA transcription. Possible candidates include chromatin remodelling factors, such as the histone chaperone FACT (FAcilitates Chromatin Transcription) or Spt6, considered to act as recycling factors for nucleosomes after the passage of RNAPII at coding regions in the euchromatin^44,45^. *Drosophila* and human Spt6 have been shown to contribute to CENP-A maintenance during interphase^31^. Thus, we tested whether these factors also contribute to Cnp1 recycling when central core ncRNAs are transcribed.

First, we determined the level of GFP-Cnp1 intensity over time during metaphase in the knockout mutant of Spt6 (*spt6*Δ) and the FACT component Pob3 (*pob3*Δ). Cells were arrested to metaphase using CBZ as well as through the *alp12-1828* mutation (**Fig. 5a**). GFP-Cnp1 intensity decreased over time in *spt6*Δ cells but not in *pob3*Δ cells. The temporal kinetics of GFP-Cnp1 levels in *pob3*Δ *spt6*Δ double-knockout cells was similar to that in *spt6*Δ cells (**Fig. 5b** and **Supplementary Fig. S4d**). This finding demonstrated that Spt6 recycles Cnp1 in WT cells, as a certain level of RNAPII leaks into the core centromere during mitosis.

**Fig. 5.**
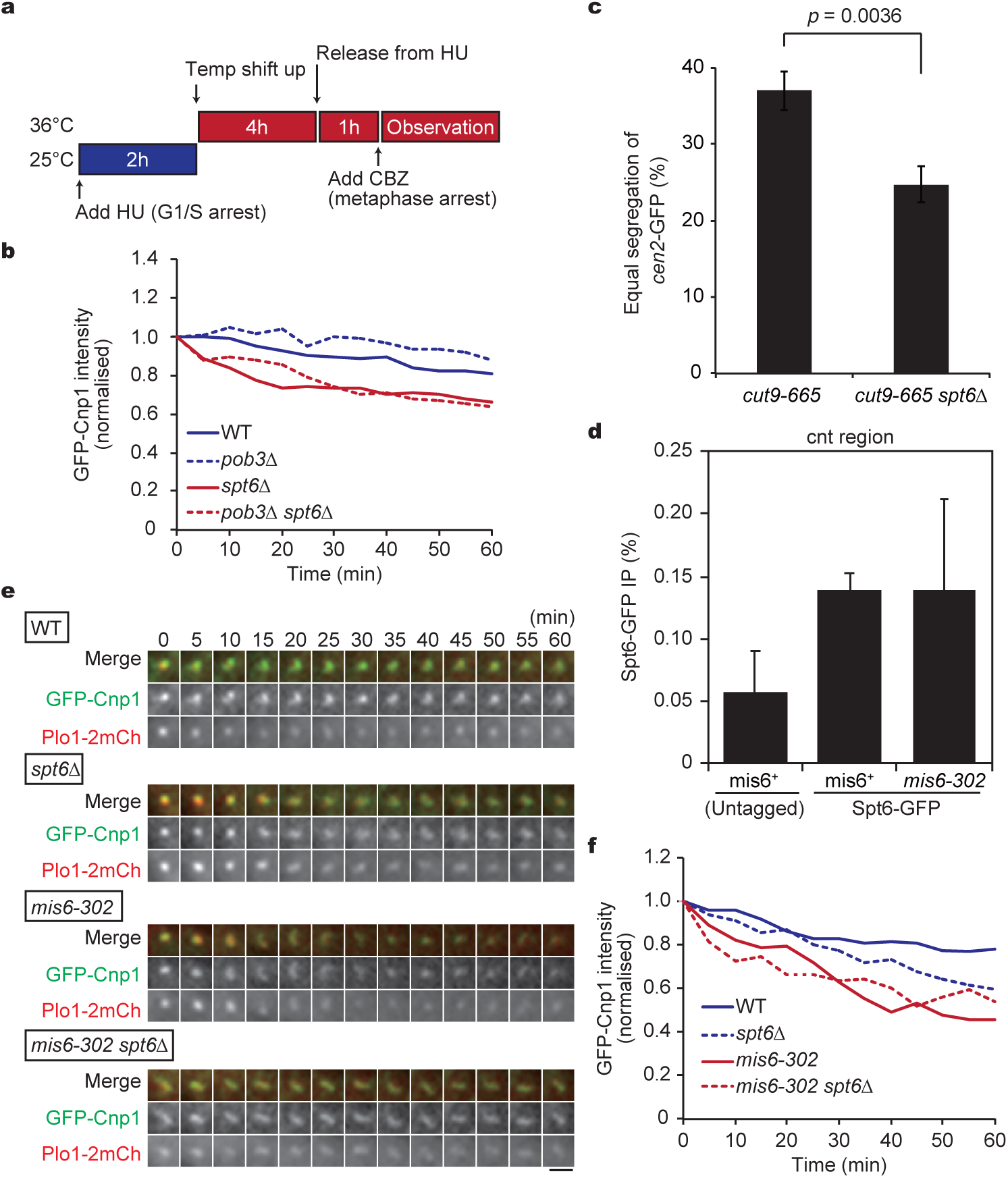
Recycling of Cnp1 requires the chromatin remodelling factor Spt6. **(a, b)** Cnp1 maintenance assays during metaphase were performed using mutants of indicated chromatin remodellers. **(a)** Experimental design. Cells arrested to G1/S at 25°C were shifted up to 36°C to inactivate Alp12. Cells were then released into mitosis and arrested at metaphase via CBZ treatment prior to observation. **(b)** A Cnp1 maintenance assay. Temporal kinetics of GFP-Cnp1 fluorescence intensity in metaphase were monitored for each mutant. WT, *n* = 18 cells; *pob3*Δ, *n* = 14 cells; *spt6*Δ, *n* = 16 cells; *pob3*Δ *spt6*Δ, *n* = 35 cells. The data are normalised to intensities at 0 min, and the mean is shown. **(c)** Population of cells with equal segregation of *cen2-*GFP (chromosome II centromere was visualised with GFP). Error bars = ± s. d., *N* = 3 experiments. *p*: Student’s t-test (two-tailed). **(d)** ChIP of Spt6-GFP using at the *cnt* region in *mis6*^+^ and *mis6-302* cells expressing Spt6-GFP or untagged Spt6. ChIP was performed with the anti-GFP antibody. Error bars = ± s. d., *N* = 3 experiments. **(e, f)** Cnp1 maintenance assay. **(e)** Time-lapse images for cells expressing GFP-Cnp1 (green) and Plo1-2mCherry (red) for each mutant background. Scale bar = 2 *µ*m. **(f)** WT, *n* = 16 cells; *mis6-302, n* = 20 cells; *spt6*Δ, *n* = 14 cells; *mis6-302 spt6*Δ, *n* = 11 cells.

During the interphase of *Drosophila* cells, HJURP deposits *de novo* CENP-A to centromeres, which is also maintained by Spt6^31^. In contrast, *de novo* CENP-A deposition does not occur during metaphase, suggesting that the role of Spt6 in CENP-A recycling may be particularly crucial in metaphase. Therefore, we investigated whether recycling Cnp1 by Spt6 during metaphase is crucial for chromosome segregation by assessing the segregation pattern of the *cen2*-GFP signal, with the centromeres of chromosome II visualised using GFP^48^. Cells were arrested to metaphase using the *cut9-665* mutant, followed by release into anaphase by re-activating Cut9 function. As shown in **Fig. 5c**, deletion of Spt6 significantly reduced the number of cells showing equal segregation of *cen2*-GFP among *cut9-665* cells.

In the absence of functional Mis6, a large amount of RNAPII appeared to surge into the central core region. A previous study indicated a direct interaction of Spt6 with RNAPII in human cells^49^. Therefore, we assessed whether there was an upregulation of Spt6 in the *mis6-302* mutant. Chromatin IP assays using Spt6-GFP pull-downs demonstrated that DNA fragments corresponding to the central core region were not specifically enriched in the *mis6-302* sample compared to the WT (*mis6*^+^) sample (**Fig. 5d**). As the amount of the histone recycler Spt6 at the central core region was unaffected despite RNAPII accumulation, Spt6 may not be able to sufficiently cope with the upsurge of ncRNA transcription occurring in the absence of Mis6.

This led us to postulate that two distinct mechanisms contribute to CENP-A maintenance during mitosis, that is, the Mis6-mediated system impedes the upsurge of RNAPII into the central core region, whereas Spt6 (FACT) recycles CENP-A in response to ncRNA transcription.

To investigate this relationship, we compared the level of Cnp1 maintenance during metaphase of *mis6-302* and *spt6*Δ cells. Although both mutants were defective in Cnp1 maintenance during metaphase, *mis6-302* cells showed more severe defects than *spt6*Δ cells (**Fig. 5e, f**). Severe reduction of the GFP-Cnp1 signal at centromeres was also detected in the *mis6-302 spt6*Δ double mutant to a similar degree as in *mis6-302* (**Fig. 5e, f**). These results indicate that once Cnp1 is lost by an upsurge of ncRNA transcription in the *mis6* mutant, the Spt6-mediated recycling system cannot fully operate for Cnp1 maintenance.

Taken together, we conclude that Mis6 is the primary factor that maintains Cnp1 during mitosis by blocking the invasion of RNAPII, which minimises the transcription of centromeric ncRNAs. The slight transcription leakage over the blockade causes the removal of Cnp1, which is then reintroduced via Spt6-mediated recycling.

## Discussion

### The life cycle of CENP-A nucleosomes

This study provides a new model for the temporal regulation of CENP-A nucleosomes during the cell cycle. That is, histone octamers containing CENP-A undergo cycles of deposition and turnover at the centromeric DNA. In fission yeast, CENP-A deposition is thought to occur during S phase (upon DNA replication) as well as during the G2 phase^7,50^. Our FRAP analysis highlighted the incorporation of CENP-A into centromeres during the G1 phase (**Fig. 1**). After photobleaching, Cnp1-GFP intensity at interphase centromeres did not recover to its original state. Assuming that the turnover of pre-existing CENP-A or H3 from centromeres may be a prerequisite for the incorporation of new CENP-A, eviction may represent a rate-limiting step for subsequent CENP-A deposition in *S. pombe*.

We propose that CENP-A can be loaded onto centromeres at any time, except during mitosis (pro ∼ metaphase). This is in line with the behaviour of the Mis18 complex and Scm3 (HJURP) throughout the cell cycle, both dispersing from kinetochores during early mitosis but returning in late mitosis (anaphase)^11,16^. In contrast, the timing of deposition is limited to the early G1 phase in human cells^8,13,14^. This discrepancy may be related to differences in the regulation of cyclin-dependent kinase (CDK) activity in each organism.

In human cells, phosphorylation of the Mis18 complex and HJURP by CDK prevents their localisation to centromeres. Since multiple distinct CDKs operate, e.g., Cdk1/2-cyclin A (active from S/G2 phase until metaphase) and Cdk1/cyclin B (pro ∼ metaphase), CENP-A deposition by the Mis18 complex and HJURP is restricted only to the G1 phase, when they escape CDK phosphorylation^51^.

In fission yeast, a single CDK (Cdc2-Cdc13/cyclin B) may determine its substrates depending on the total level of activity^52^. It is possible that the Mis18 complex and HJURP are phosphorylated only when CDK activity is substantially high, that is, during metaphase. This might explain why fission yeast cells are competent in CENP-A deposition at any time except metaphase. As deposition machinery is compromised during metaphase, cells employ specific mechanisms to not lose CENP-A nucleosomes meanwhile.

### The mechanism of CENP-A maintenance by CENP-I

The present study elucidates a previously unexplored mechanism through which CENP-A is maintained at the central core region of centromeres. In particular, we demonstrated a new role for Mis6 (CENP-I) as a CENP-A maintenance factor. Mis6 has been considered a factor required for CENP-A deposition, since the localisation of Cnp1 and Scm3 is lost in the *mis6* mutant^15,32^. However, we found that in the absence of functional Mis6, CENP-A at centromeres decreased during metaphase when CENP-A was no longer deposited. Therefore, this reduction directly reflects CENP-A turnover from the centromeres.

A recent study revealed that HJURP, in cooperation with the MCM complex, is required for CENP-A maintenance during S phase in human cells^30^. However, we demonstrated that CENP-A was maintained from G1/S phase until the end of mitosis in the *scm3* mutant (**Fig. 2c**). This suggests that Scm3 (HJURP) does not predominantly contribute to CENP-A maintenance in fission yeast. Instead, it is possible that Mis6 maintains CENP-A during interphase in addition to metaphase.

Transcription of centromeric ncRNA was previously detected throughout the cell cycle in *S. pombe*^47^, with only a certain fraction of RNAPII localising into the central core (*cnt*) region to transcribe ncRNAs^23^. As our results demonstrated that RNAPII binding to the *cnt* region was elevated in the *mis6* mutant, we propose that Mis6 (CENP-I) can act as an insulator for RNAPII invasion into the central core region in other organisms as well. In human cells, unknown factors in addition to HJURP appear to maintain CENP-A^30^. Moreover, ncRNA transcription at centromeres was detected in higher eukaryotes besides humans, including mice and the tammar wallaby^21^, suggesting that CENP-I possibly maintains CENP-A in these species as well. On the other hand, *CENP-I-* deficient chicken DT40 cells display stable localisation of CENP-A to centromeres^53,54^, suggesting that CENP-I might not contribute to CENP-A maintenance. Assuming that CENP-I serves as an insulator, it would be intriguing to investigate whether ncRNAs are actively transcribed at centromeres in DT40 cells. Molecular schemes for CENP-A maintenance may be intimately linked to whether ncRNA transcription occurs in centromeres. The utilisation of Mis6 (CENP-I) represents a suitable solution in organisms which actively transcribe centromeric ncRNAs, as it contributes to both CENP-A deposition and CENP-A maintenance as an RNAPII insulator.

### Two-step machinery for sustainable CENP-A positioning during mitosis

Even though Mis6 impedes the progression of RNAPII into the *cnt* region of centromeres, transcripts are still detected at a certain level^23^. This indicates that CENP-A in the *cnt* region is temporarily removed by the chromatin remodelling factor Fft3 when RNAPII proceeds but is mostly maintained by the histone recycler Spt6. Despite the accumulation of RNAPII within the *cnt* region of the *mis6* mutant, concomitant accumulation of the histone recycler Spt6 was not observed. Unlike in human cells^49^, *S. pombe* Spt6 might not directly interact with RNAPII. Alternatively, an intense upregulation of RNAPII in the *cnt* failed to efficiently recruit Spt6. In either case, the capacity of Spt6 at the central core region appears to be limited to recycling CENP-A only at basal levels. We conclude that two mechanisms operate for CENP-A maintenance. First,

Mis6 serves to insulate against the invasion of RNAPII into the core centromere, thus maintaining CENP-A, particularly during mitosis when *de novo* CENP-A deposition does not occur. Although Spt6 recycles CENP-A during mitosis in WT cells as the second machinery for CENP-A maintenance, the recycling capacity of Spt6 appears to be limited. Insulation of RNAPII by Mis6 is thus employed as the primary machinery, followed by Spt6-mediated CENP-A recycling as a backup, in a stepwise strategy for the epigenetic maintenance of CENP-A until the next cell cycle.

## Acknowledgements

We thank Y. Takayama, A. Yamamoto, Y. Hiraoka, T. Sakuno Y. Watanabe, M. Yanagida and the National Bioresource Project (NBRP) of Japan for the yeast strains. H.H. was a research fellow of the Japan Society for the Promotion of Science (JSPS; 16J09035). This study was supported by JSPS KAKENHI JP25291041, JP15H01359, JP16H04787, JP16H01317, JP18K19347 and 21H00261 to M.S. This study was also supported by The Uehara Memorial Foundation, Ohsumi Frontier Science Foundation and Waseda University grants for Special Research Projects 2017B-242, 2017B-243, 2018B-222, 2019C-570 and 2020R-038 to M.S. This work was also partly supported by the JSPS Core-to-Core Program, A: Advanced Research Networks.

## Author Contributions

H.H. and M.S. conceived and designed the experiments. H.H. and Y.S. performed the experiments. H.H. wrote the draft of the manuscript. M.S. supervised the project and finalised the manuscript through discussion with H.H. and Y.S.

## Methods

### *S. pombe* strains and genetics

Strains used in this study are listed in **Supplementary Table S1**. Standard PCR-based methods for gene targeting were employed for the construction of knock-out mutants and strains with fluorescent protein tagging^55-57^. We used multiple constructs for the visualisation of Cnp1. In **Figure 1**, strains expressing the Cnp1-GFP gene under an *adh21* promoter at the *C* locus (adjacent to the SPAC26F1.12c gene of chromosome 1) were used (the original strain was a gift from Y. Watanabe)^58^. In **Fig. 2C**, GFP-Cnp1 was expressed under the endogenous promoter (a gift from Y. Takayama)^59^. In other figures, the fusion gene of GFP-Cnp1 or mCherry-Cnp1 driven by the *nmt1* promoter was inserted at the *CO2* locus of chromosome 2 (adjacent to the SPBPB7E8.01 gene)^60^ as an extra copy of the endogenous *cnp1*^+^ gene.

To create these integrant strains, we first created plasmids harboured the *nmt1* promoter placed upstream of the *cnp*1 coding sequence fused with the GFP or mCherry gene via the Golden Gate method^60^. The constructed plasmids were digested by the FseI enzyme for linearisation and introduced into strains to induce homologous recombination at the *CO2* locus.

For knocksideways experiments, expression of Kis1-GBP and Kis1-GBP-mCherry was induced via pREP1-based plasmids containing the *nmt1* promoter. The *GFP-kan* gene was inserted to the end of the *mis6*^+^ coding sequence, so that the strain expresses the fusion protein of Mis6-GFP instead of the endogenous Mis6 protein.

Cells in **Figs. 1, 2, 4, 5, S1, S3** and **S4** were cultured in YE5S medium, while those in **Figs. 3** and **S2** were cultured in Edinburgh minimal medium (EMM).

### Mutagenesis of the *scm3* gen

We first constructed the *scm3-myc-hph* strain, in which the *scm3*^+^ gene was tagged with the c-Myc epitope at the C-terminus marked with the *hph* selection marker gene, which confers resistance to 100 *µ*g ml^-1^ hygromycin B. We then induced random point mutations into *scm3-myc-hph* DNA fragments through an error-prone PCR reaction. The mutated fragments were then introduced into the *mis6-GFP-kan* strain to induce homologous recombination with the endogenous *scm3*^+^ gene. Colonies grown on YE5S + hygromycin B at 25°C were replicated onto YE5S plates containing 2 *µ*g ml^-1^ phloxine B at 36°C. Colonies showing temperature sensitivity with the normal localisation of Mis6-GFP were selected for sequencing and further experiments.

### Chromatin IP

Cells were cultured at 25°C in YE5S, followed by a temperature shift up to 36°C for 6 h. The cells were then fixed with 1% formaldehyde for 10 min at 36°C and left on ice for 50 min. Chromatin IP assays in this study were carried out as previously described^11^, with minor modifications. In brief, fixed cells were washed four times with Buffer I (50 mM HEPES/NaOH [pH 7.5], 140 mM NaCl, 1 mM EDTA [pH7.5], 1% Triton X-100 and 0.1% sodium deoxycholate) at 4°C and kept frozen at –80°C. Cells were then suspended in Buffer I supplemented with Complete Protease Inhibitor cocktail (Roche) and 1 mM phenylmethylsulfonyl fluoride. Cells were destroyed using acid-washed glass beads and a FastPrep-24 bead shocker (4 × 20 sec, power = 6.0). The cell lysates were then sonicated using a sonifier VP-050 (PWM 10%). A round of iterative sonication for 10 sec comprising ON (0.2 sec) and OFF (0.4 sec) was repeated 10 or 15 times to shear chromatin DNA into fragments. Lysates were then centrifuged (14,000 rpm for 15min at 4°C) to collect supernatants, and the concentration was adjusted to 10 mg ml^-1^.

For immunoprecipitation, the rabbit anti-RNA polymerase II (phosphoS5) polyclonal antibody (1:100; ab5131) or rabbit anti-GFP polyclonal antibody (1:250; Clontech, 632592) was incubated with 200 µL of the lysate for 1 h at 4°C. Protein A sepharose (GE) was then added, incubated for 2 h at 4°C and washed three times with Buffer I, followed by suspension with Buffer I’ (50 mM HEPES/NaOH [pH 7.5], 500 mM NaCl, 1 mM EDTA [pH 7.5], 1% Triton X-100 and 0.1% sodium deoxycholate) thrice, Buffer II (10 mM Tris-HCl [pH 8.0], 250 mM LiCl, 0.5% NP-40, 0.5% sodium deoxycholate) twice and TE twice. The amounts of DNA derived from whole-cell extracts and ChIP samples were assessed via quantitative PCR on the StepOne Real-Time PCR system (Applied Biosystems) using SYBR Green (TOYOBO). Oligonucleotide primers for detecting the central core region of centromeres were used as previously described^61^.

### Microscopy

Living cells were transferred to a glass-bottom dish (Matsunami) pre-coated with lectin, and the dish was filled with liquid EMM preheated at 25°C or 36°C. Mounted cells were observed at 25°C or 36°C using a microscope (IX71, Olympus) with the DeltaVision-SoftWoRx system (Applied Precision), as previously described^62^. Cells were imaged in 12 sections at 0.4-*µ*m intervals along the z-axis. The temperature conditions are shown below. In **Fig. 1**, Cells were observed for 1 h after culturing at 36°C for 3 h. In **Fig. S1d**, overnight cultures were incubated at 36°C for 4 h, and cells were fixed with 3.2% formaldehyde (Thermo Fisher Scientific) for 20 min. In **Figs. 3c** and **S2**, cells were cultured at 25°C for 20 h in EMM and imaged. The acquired images were processed as follows: images taken along the z-axis were deconvoluted and projected into a single image using the Quick Projection algorithm in the SoftWoRx software (v3.7.0 and v.6.5.1). The fluorescence intensity of Cnp1 visualised with GFP or mCherry was measured using the Data Inspector command in SoftWoRx. The mean intensity of a single dot signal of Cnp1-GFP or GFP-Cnp1 in 6 × 6 pixels was measured, and the background signal outside of the nucleus was subtracted. For photobleaching of a Cnp1-GFP dot signal, the dot area was successively irradiated four times by the 488 nm laser (50% in power; Seki Technotron) for 0.05 sec using the SoftWoRx-QLM system. In **Figs. 1, 2, 3c, 4, 5** and **S4**, the resolution of enlarged images was adjusted from 72 to 144 pixels inch^-1^ using Adobe Photoshop (ver. 2021).

### Drug treatment

To inhibit RNAPII, 1,10-phenanthroline (Sigma-Aldrich) diluted to 100 mg ml^-1^ with ethanol was added to the EMM at a final concentration of 100 *µ*g ml^-1^. Alternatively, thiolutin (Wako) diluted to 5 mg ml^-1^ with DMSO was added to EMM at a final concentration of 20 *µ*g ml^-1^ as previously described^63^. For mock treatment, the same amount of solvent (ethanol or DMSO) was added to the EMM. Observations began 15 min after the addition of reagents. To arrest cells at metaphase for more than 1 h, the microtubule poison carbendazim (CBZ; Sigma-Aldrich) diluted to 5 mg ml^-1^ with DMSO was added to EMM at a final concentration of 50 *µ*g ml^-1^ immediately before observation. To arrest cells during the G1/S phase, hydroxyurea (HU; Sigma-Aldrich) diluted to 1.5 M with DMSO was added to YE5S at a final concentration of 15 mM.

### Chromosome segregation assay with *cen2-*GFP

To evaluate chromosome segregation in *cut9-665* and *cut9-665 spt6Δ* mutants, the *cen2-*GFP system was employed to visualise the position of chromosome II centromere with GFP^48^. For metaphase arrest, cells were first arrested at the G1/S phase using HU for 2 h. In the presence of HU, the temperature was then shifted up to 36°C to inactivate Cut9 (an APC/C component). Cells were then washed with ddH^2^O three times and cultured in YE5S for 2 h at 36°C to release cells from G1/S arrest until metaphase arrest via Cut9-inactivation. The culture was shifted down to 25°C again to finally release cells from metaphase to anaphase, and the distribution of *cen2*-GFP dots was observed for 1 h.

