## Supplementary Information for "Mis6/CENP-I maintains CENP-A nucleosomes against centromeric non-coding transcription during mitosis"

Supplementary table S1

Supplementary figures S1–S4

### Supplementary table S1. *S. pombe* strains used in this study

| Strain No. | Genotype | Figures |
|---|---|---|
| HH609 | *$h^{+}$ cdc10-27 C::Padh21-cnp1-GFP-kan cnp3-tdtomato-hph sid4-CFP-nat leu1* | 1a, b |
| HH614 | *$h^{90}$ cdc25-22 C::Padh21-cnp1-GFP-kan cnp3-tdtomato-hph sid4-CFP-nat* | 1c, d |
| HH1521 | *$h^{-}$ alp12-1828 C::Padh21-cnp1-GFP-kan plo1-2mCherry-hph sid4-CFP-nat leu1 ura4 ade6-M216* | 1e, f |
| SP1769 | *$h^{-}$ GFP-cnp1-nat leu1 ura4 lys1* | 2a |
| HH1812 | *$h^{+}$ GFP-cnp1-nat mis6-302-HA-bsd leu1 ura4 ade6-M216* | 2a |
| HH1928 | *$h^{+}$ cut9-665 GFP-cnp1-nat leu1 ade6-M216* | 2a |
| HH1930 | *$h^{+}$ cut9-665 GFP-cnp1-nat mis6-302-HA-bsd leu1 ura4 ade6-M216* | 2a |
| HH1888 | *$h^{+}$ GFP-cnp1-nat ndc80-mCherry-bsd leu1 ura4 ade6-M216* | 2c |
| HH1890 | *$h^{+}$ $scm3^{S55P}$-13myc-hph GFP-cnp1-nat ndc80-mCherry-bsd leu1 ura4 ade6-M216* | 2c |
| HH1897 | *$h^{-}$ mis6-302 GFP-cnp1-nat ndc80-mCherry-bsd leu1 ura4 ade6-M216* | 2c |
| HH1687 | *$h^{+}$ alp12-1828 CO2::nmt1-GFP-L-cnp1-kan plo1-2mCherry-hph sid4-CFP-nat leu1 ura4 ade6-M210* | 2e, f, g, h, 4d, e 5b, e, f, S4 |
| HH1693 | *$h^{-}$ alp12-1828 CO2::nmt1-GFP-L-cnp1-kan plo1-2mCherry-hph sid4-CFP-nat mis6-302-HA-bsd leu1 ura4 ade6-M216* | 2e. f, 4d, e, 5e, f S4a, b, c |
| SG653 | *$h^{-}$ alp12-1828 CO2::nmt1-GFP-L-cnp1-kan plo1-2mCherry-hph sid4-CFP-nat mis12-537-3HA-bsd leu1 ura4 ade6-M216* | 2g |
| SG442 | *$h^{90}$ alp12-1828 CO2::nmt1-GFP-L-cnp1-kan plo1-2mCherry-hph sid4-CFP-nat nuf2-2::$ura4^{+}$ leu1 ura4 ade6-M210* | 2h |
| HH503 | *$h^{+}$ mis6-GFP-kan Z2-CFP-atb2-nat leu1 ura4 his2 ade6-M216* +pREP1-*kis1-GBP-mCherry* | 3b |
| HH1678 | *$h^{-}$ mis6-GFP-kan CO2::nmt1-mCherry-L-cnp1-hph Z2-CFP-atb2-nat leu1 ura4 ade6-M216* +pREP1-*kis1-GBP* | 3c, d, S2 |
| HH1684 | *$h^{+}$ mis6-GFP-kan CO2::nmt1-mCherry-L-cnp1-hph Z2-CFP-atb2-nat leu1 ura4 his2 ade6-M216* +pREP1 | 3c, d, S2 |
| HH333 | *$h^{-}$ leu1 ura4 ade6-M216* +pREP1 | 3e |
| HH337 | *$h^{-}$ leu1 ura4 ade6-M216* +pREP1-*kis1-GBP-mCherry* | 3e |
| HH451 | *$h^{-}$ mis6-GFP-kan leu1 ura4 ade6-M216* +pREP1-*kis1-GBP-mCh* | 3e |
| HH1607 | *$h^{-}$ mis6-GFP-kan leu1 ura4 ade6-M216* +pREP1 | 3e |
| FY336 | *$h^{-}$ cnt1::$ura4^{+}$ leu1 ura4-DS/E ade6-M210* | 4a, S3a |
| HH2136 | *$h^{-}$ mis6-302 cnt1::$ura4^{+}$ leu1 ura4-DS/E ade6-M210* | 4a, S3a |
| JY741 | *$h^{-}$ leu1 ura4 ade6-M216* | 4b, 5d, S3b |
| KA2130 | *$h^{90}$ mis6-302 leu1 ura4 ade6-M216* | 4b, S3b |
| HH2001 | *$h^{-}$ alp12-1828 CO2::nmt1-GFP-L-cnp1-kan plo1-2mCherry-hph sid4-CFP-nat pob3::bsd leu1 ura4 ade6-M216* | 5b, S4d |
| HH2010 | *$h^{+}$ alp12-1828 CO2::nmt1-GFP-L-cnp1-kan plo1-2mCherry-hph sid4-CFP-nat spt6::$ura4^{+}$ leu1 ura4 ade6-M210* | 5b, S4d |
| HH2029 | *$h^{+}$ alp12-1828 CO2::nmt1-GFP-L-cnp1-kan plo1-2mCherry-hph sid4-CFP-nat pob3::bsd spt6::$ura4^{+}$ leu1 ura4 ade6-M210* | 5b, S4d |

| | | |
|---|---|---|
| HH2061 | $h^{90}$ cut9-665 cen2<<lac0-kan-ura4+ his7+<<(dis1pro)-GFP-lacI spt6::$ura4^{+}$ leu1 ura4 his2 ade6-M216 | 5c |
| HH2066 | $h^{90}$ cut9-665 cen2<<lac0-kan-ura4+ his7+<<(dis1pro)-GFP-lacI leu1 ura4 his2 ade6-M216 | 5c |
| HH1971 | $h^{-}$ mis6-302 spt6-GFP-kan leu1 ura4 ade6-M210 | 5d |
| HH1975 | $h^{-}$ spt6-GFP-kan leu1 ade6-M216 | 5d |
| HH2009 | $h^{-}$ alp12-1828 CO2::nmt1-GFP-L-cnp1-kan plo1-2mCherry-hph sid4-CFP-nat spt6::$ura4^{+}$ leu1 ura4 ade6-M216 | 5e, f |
| HH2055 | $h^{+or-}$ alp12-1828 CO2::nmt1-GFP-L-cnp1-kan plo1-2mCherry-hph sid4-CFP-nat mis6-302-HA-bsd spt6::$ura4^{+}$ leu1 ura4 ade6-M216 | 5e, f |
| HH322 | $h^{-}$ mis6-GFP-kan leu1 ura4 ade6-M216 | S1a, b, c |
| HH1820 | $h^{-}$ mis6-GFP-kan $scm3^{S55P}$-13myc-hph leu1 ura4 ade6-M216 | S1a, b, c |
| KRY211 | $h^{+}$ mis6-2GFP-kan cnp3-tdtomato-hph sid4-CFP-nat leu1 ura4 ade6-M216 | S1d |
| SG366 | $h^{+}$ mis12-537 mis6-2GFP-kan cnp3-tdtomato-hph sid4-CFP-nat leu1 ura4 his2 ade6-M216 | S1d |
| SG342 | $h^{+}$ nuf2-1::$ura4^{+}$ mis6-2GFP-kan cnp3-tdtomato-hph sid4-CFP-nat leu1 ura4 his2 ade6-M216 | S1d |
| SG344 | $h^{+}$ nuf2-2::$ura4^{+}$ mis6-2GFP-kan cnp3-tdtomato-hph sid4-CFP-nat leu1 ura4 his2 ade6-M216 | S1d |
| SG571 | $h^{-}$ mis12-537 cnt1::$ura4^{+}$ leu1 ura4-D18 or ura4-DS/E ade6-M210 | S3a |
| SG594 | $h^{-}$ nuf2-2::$ura4^{-}$ cnt1::$ura4^{+}$ leu1 ura4-D18 or ura4-DS/E ade6-M210 | S3a |
| MT111 | $h^{+}$ mis12-537 leu1 ura4 his2 ade6-M216 | S3b |
| YK2330 | $h^{-}$ nuf2-1::$ura4^{+}$ leu1 ura4 ade6-M210 | S3b |
| HH2000 | h- alp12-1828 CO2::nmt1-GFP-L-cnp1-kan plo1-2mCherry-hph sid4-CFP-nat mis6-302-HA-bsd fft3::$ura4^{+}$ leu1 ura4 ade6-M216 | S4a |

SP1769 is a gift from Y. Takayama, from which HH1812, HH1888, HH1890, HH1897, HH1928, and HH1930 were created. Original strains used for creation of YK1316, YK2330, HH2061, HH2066, SG342, SG344, SG442 and SG594 were gifted from Y. Hiraoka and A. Yamamoto. The original strain used for creation of HH609, HH614, and HH1521 were gifted from Y. Watanabe and T. Sakuno. Original strains used for creation of HH1521, HH1687, HH1693, HH1928, HH1930, HH2000, HH2009, HH2010, HH2055, HH2061, HH2066, SG442 and SG653 were gifted from T. Toda. Original strains used for creation of KA2130, MT111, HH1693, HH1812, HH1897, HH1930, HH2000, HH2136, SG366, SG571 and SG653 were gifted from M. Yanagida. FY336 strain was provided from NBRP (originated from M. Yanagida), from which HH2136, SG571 and SG594 were made. Other strains are our stock.

### Supplementary figures

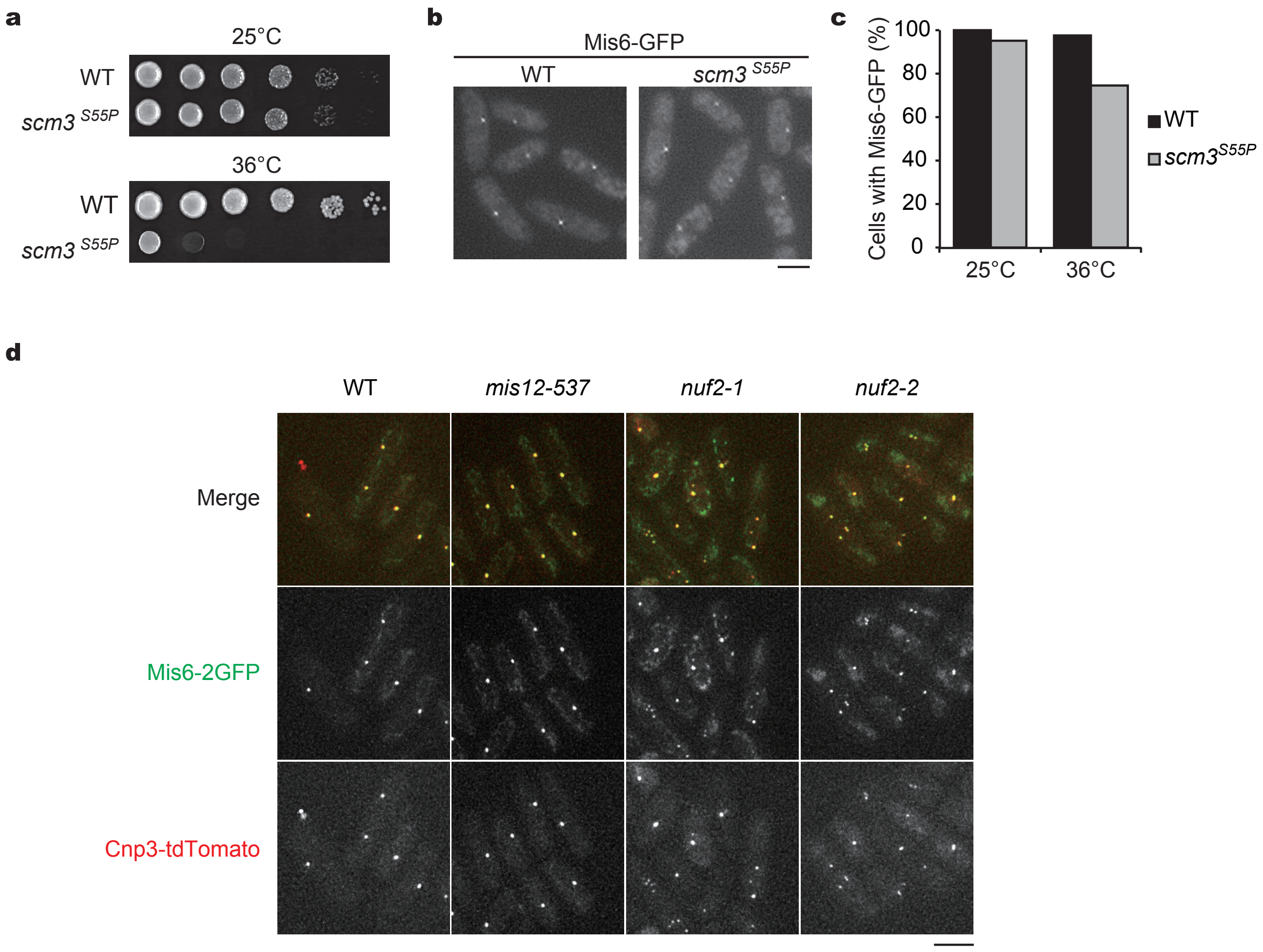


**Supplementary figure S1 Mis6 localises properly in various kinetochore mutants**
**(a)** Growth assays for indicated strains. Ten-fold serial dilutions of WT and *scm3*$^{S55P}$ cells were grown at 25˚C or 36˚C. **(b)** Mis6-GFP localised normally to centromeres in *scm3*$^{S55P}$ cells as in WT cells. Bar = 5 $\mu$m. **(c)** Percentages of cells with Mis6-GFP at centromeres in WT and the *scm3*$^{S55P}$ cells at 25°C and 36°C (6 h) ($n > 130$ cells). **(d)** Mis6-2GFP (green) colocalised with kinetochore marker (Cnp3-tdTomato, red) in *mis12-537*, *nuf2-1* and *nuf2-2* cells as with WT cells at 36°C (4h) ($n > 93$ cells). Bar = 5 $\mu$m.

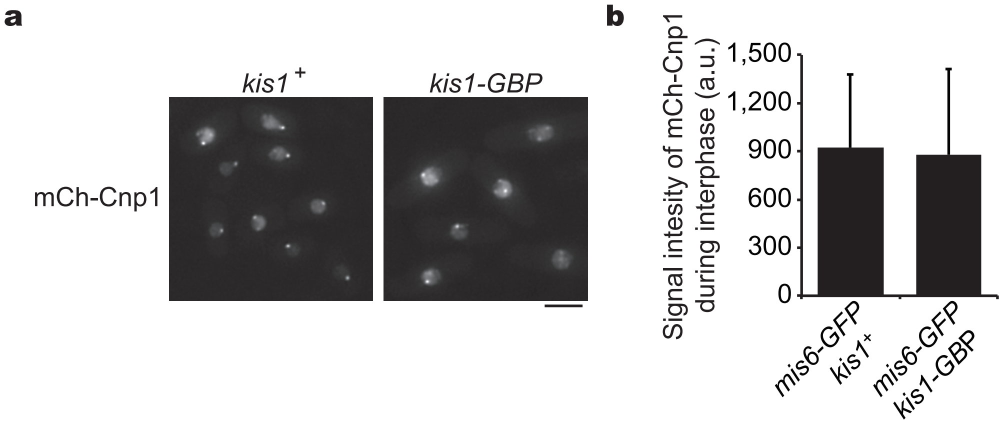


**Supplementary figure S2 Interphase localisation of Cnp1 is not affected in the *mis6-GFP kis1-GBP* strain**

**(a)** mCherry-Cnp1 localised normally to centromeres in interphase cells of *kis1*$^{+}$ *mis6-GFP* and *kis1-GBP mis6-GFP* strains. Bar = 5 $\mu$m. **(b)** Fluorescence intensities of mCherry-Cnp1 at centromeres in interphase were quantified in both strains ($n > 50$ cells, mean ± s. d.).

**a**

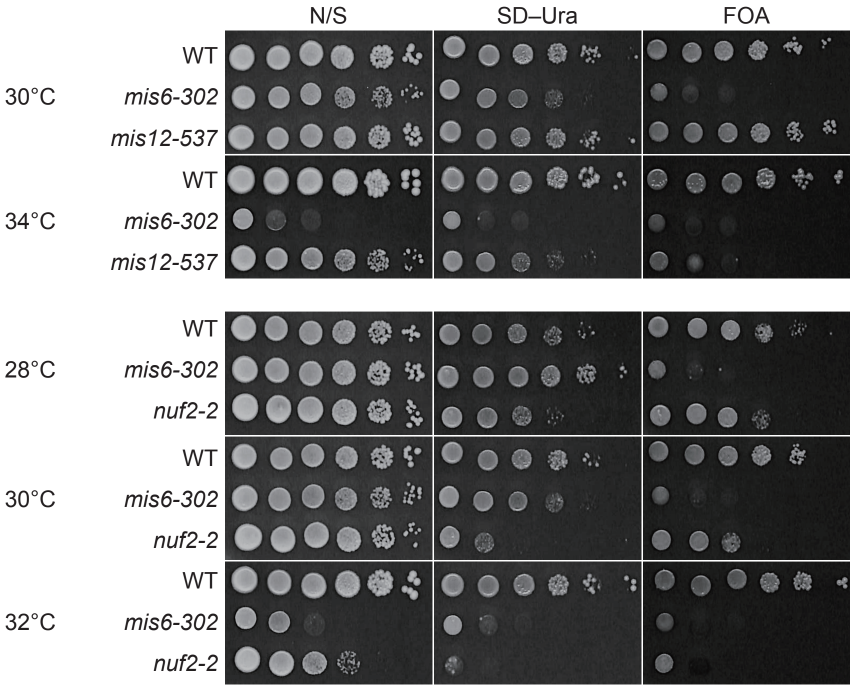


**b**

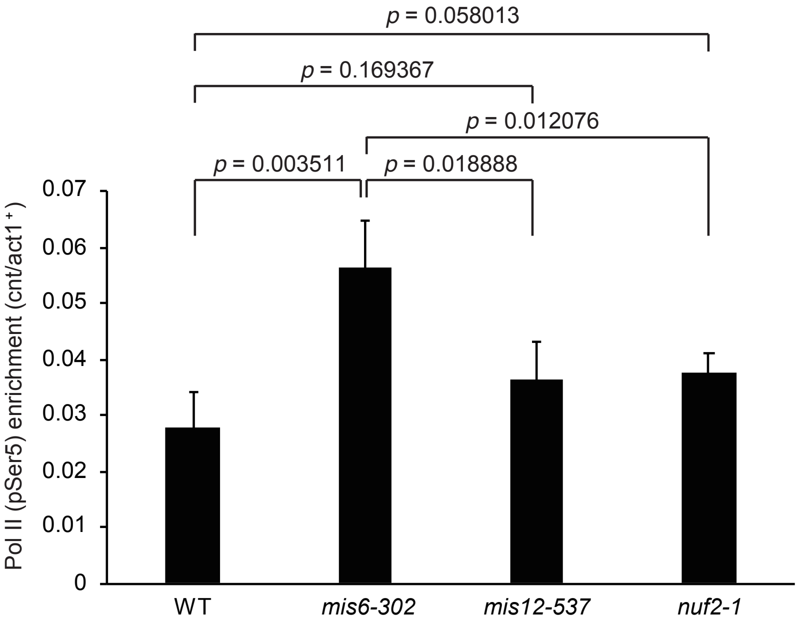


**Supplementary figure S3 Silencing at the core centromere is intact in the *mis12* and *nuf2* mutant cells**

**(a)** Silencing assays. Ten-fold serial dilution of WT, *mis6-302*, *mis12-537* and *nuf2-2* cells with *cnt1::ura4+*, grown on N/S, SD-Ura and FOA media at indicated temperature. **(b)** ChIP assays for RNAPII pSer5 in WT, *mis6-302*, *mis12-537* and *nuf2-1* cells at 36˚C (4h). The % input of the *cnt1* region was normalised to the % input of the *act1+* region. Error bars = ± s. d., *N* = 4 experiments. *p*: Student's t-test (two-tailed).

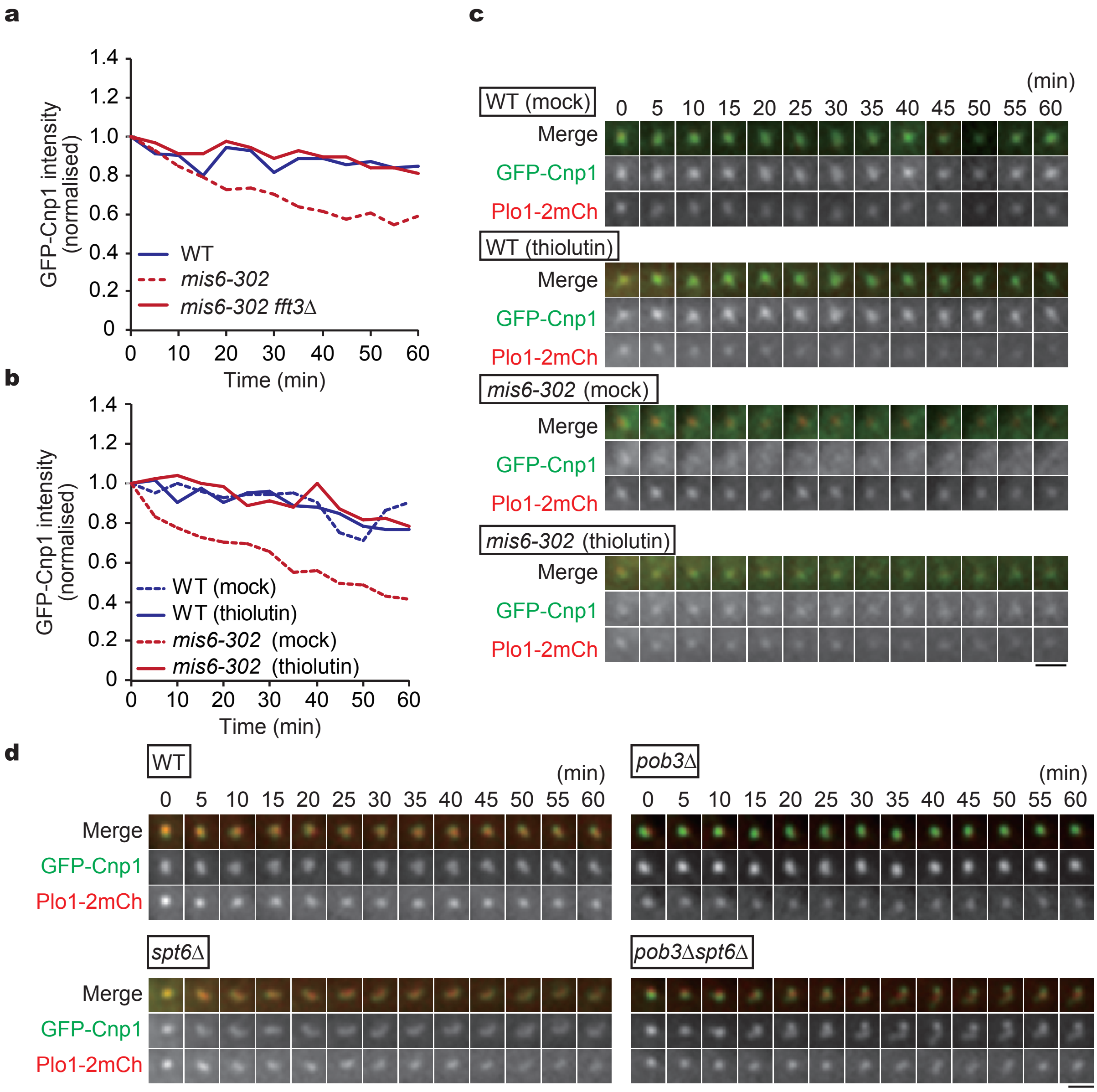


**Supplementary figure S4 Assays for mitotic Cnp1 maintenance in various conditions**
**(a)** Temporal kinetics of the GFP-Cnp1 intensity in pro ~ metaphase of indicated cells. WT, *n* = 15 cells; *mis6-302*, *n* = 18; *mis6-302 fft3*Δ, *n* = 13. **(b, c)** Temporal kinetics of the GFP-Cnp1 intensity in pro ~ metaphase of WT and *mis6-302* cells in the presence of thiolutin. WT (mock), *n* = 23 cells; *mis6-302* (mock), *n* = 15; WT (thiolutin), *n* = 13; *mis6-302* (thiolutin), *n* = 11. The data have been normalised to intensities at 0 min. Mean is shown in the graph. Representative time-lapse images for cells expressing GFP-Cnp1 (green) and Plo1-2mCherry (red) in each genetic background. **(d)** Representative time-lapse images used for data presentation in **Figure 5b**. Bars = 2 $\mu$m.
